# Plague-driven selection enriched familial Mediterranean fever mutations in Armenia

**DOI:** 10.64898/2026.09.10.750633

**Authors:** Anahit Hovhannisyan, Mariya Antonosyan, Arsen Bobokhyan, Hasmik Simonyan, Levon Aghikyan, Pavel Avetisyan, Mikayel Badalyan, Hakob Simonyan, Meri Safaryan, Artak Gnuni, Ashot Piliposyan, Avetis Grigoryan, Meline Simonyan, Artavazd Zaqyan, Mkrtich Zardaryan, Hovann Simonian, Levon Mkrtchyan, Iseult Jackson, Emily Breslin, Bruno Ariano, Valeria Mattiangeli, Zaruhi Khachatryan, Lara M Cassidy, Levon Yepiskoposyan, Daniel G Bradley, Andrea Manica

**Affiliations:** Smurfit Institute of Genetics, Trinity College Dublin, Ireland; Department of Coevolution of Land Use and Urbanisation, Max Planck Institute of Geoanthropology, Germany; Institute of Archaeology and Ethnography, Armenia; Chair of Archaeology and Ethnography, Yerevan State University, Armenia; Service for the Protection of Historical Environment and Historical Cultural Museum Reservations, Ministry of Education, Science, Culture and Sports, Armenia; Institute of History, Armenia; Scientific Research Centre of the Historical and Cultural Heritage, Ministry of Education, Science, Culture and Sports, Armenia; Armenian DNA project at Family Tree DNA, Houston, Texas, USA; Shirak Center for Armenian Studies, National Academy of Sciences of the Republic of Armenia, Gyumri, Armenia; Institute of Molecular Biology, Armenia; Department of Zoology, University of Cambridge, UK

## Abstract

The discovery that positive selection due to resistance to malarial infection maintained the prevalence of sickle cell and other disease alleles was a landmark in evolution, genetics and epidemiology. Mutations in the *MEFV* gene causing familial Mediterranean fever with high frequency in Eastern Mediterranean populations give the only other strong candidates for a similar process. Here we estimate a pathogenic *MEFV* mutation burden in Armenians with a carrier rate of ~41%: a worldwide maximum. By combining these data with regional ancient *MEFV* genotypes spanning more than 60 centuries, we establish that the allele frequencies of three mutations M694V, M680I, and V726A underwent recent rapid increases, indicating positive selection with an onset consistent with the first *Yersinia pestis* plague pandemic, ~541 CE to 767 CE or even earlier in time, echoing functional data implying resistance to this infection as the selective agent. This extends to three the number of *MEFV* variants for which there is evidence for strong concurrent selection at this locus in the Eastern Mediterranean, making it unlikely that this geographic peak resulted from chance alone. Although bubonic plague was extremely widespread, the restricted geography of this selection points toward the Eastern Mediterranean as a longstanding enzootic centre with continuous recycling of outbreaks.

## INTRODUCTION

Familial Mediterranean fever (FMF) is the most common monogenic autoinflammatory disease worldwide (*1*). It is caused by pathogenic mutations in the *MEFV* gene (16p13.3), which encodes pyrin, a key regulator of the innate immune system (*2, 3*). Mutant pyrin forms an aberrant inflammasome complex, leading to excessive production of proinflammatory cytokines (mainly IL-1β) which drives recurrent episodes of fever, accompanied by serositis, arthritis, and, in untreated cases, secondary amyloidosis (*4, 5*). The severity of FMF is highly correlated with specific variants and, although the disease is primarily inherited in an autosomal recessive manner, studies suggest a more complex inheritance pattern, with compound heterozygotes and even single heterozygotes for certain mutations also manifesting the disease (*6*). More than 30 mutations have been associated with FMF (*7*): among these, the four most frequently observed in patients are M694V, M680I, and V726A in exon 10, and E148Q in exon 2. The disease predominantly occurs among Eastern Mediterranean groups with reported carrier rates of these four *MEFV* mutations ranging from 10-39% in e.g. Turkish, Arabic, and Jewish populations (*8, 9*) and a recent whole-genome study of the Armenian population revealed a markedly high number of individuals carrying at least one of these mutations (47%), suggesting that clinically routine FMF testing for 12 pathogenic *MEFV* mutations (so called strip-array testing, based on reverse-hybridisation to allele-specific oligonucleotide probes immobilised as an array) may substantially underestimate true disease prevalence (*10*).

This strikingly high regional frequency of *MEFV* mutations has prompted hypotheses of evolutionary advantage. It has been proposed that heterozygote carriers have increased protection against presumptive endemic disease(s) such as tuberculosis, smallpox and cholera (*11*). However, more recently, it has been persuasively argued that *MEFV* mutations can confer enhanced resilience against bubonic plague (*12*). This latter study reported evidence of positive selection of the M694V and V726A mutations from their analysis of Turkish population genomic data and suggested an underlying molecular mechanism. Their functional experiments showed that mutant pyrin binds less avidly to the *Yersinia pestis* virulence factor YopM, attenuating YopM-induced IL-1β suppression, and thereby enabling a heightened inflammatory response upon infection. However, their genetic analysis relied exclusively on a modern dataset and a comprehensive investigation incorporating ancient DNA (aDNA) from the Eastern Mediterranean region remains lacking.

Here, we conduct the first time-series based test for selection acting on *MEFV* mutations using data from Armenia, a country with strong population genetic continuity stretching over 6,000 years (*10*) and the highest prevalence of *MEFV* mutations (*10, 13*). We applied a targeted gene capture strategy to 85 ancient samples (Fig. 1A, Data S1), enriching for the *MEFV* locus and enabling reliable variant detection across a temporal transect of regional genomes extending over six thousand years beginning from the Chalcolithic. We also sequenced *MEFV* genes from 146 representatives of the modern Armenian population and combined them with 36 previously published modern samples (Data S2). Through investigation of allele-frequency trajectories and haplotype analyses we show that strong positive selection increased the frequency of three pathogenic *MEFV* variants in the Armenian population, likely around the onset of the first plague pandemic in the region.

**Figure 1.**
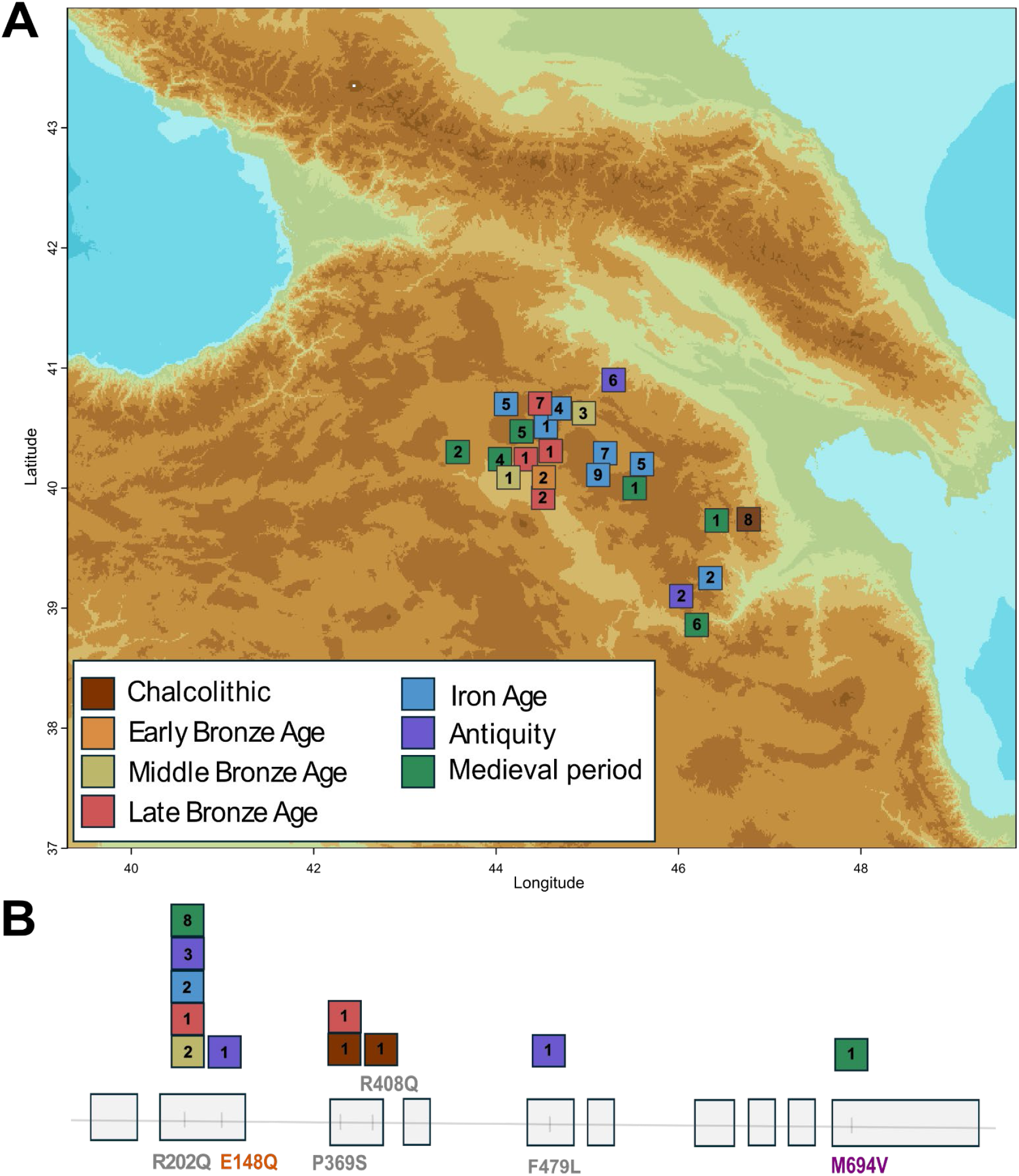
Location of *MEFV*-captured ancient samples and the incidences of gene mutation among them. **(A)** Map showing the geographic locations of ancient samples captured for the MEFV gene in this study. The legend indicates the color coding for chronological periods. Numbers inside squares represent the number of samples from each archaeological site. **(B)** MEFV mutations found among the ancient samples and exonic position of these. Color coding and numbers as above.

## RESULTS

### *MEFV* gene mutation prevalence in modern Armenian and ancient populations

We first examined 182 modern genomes (146 newly generated here) from Armenia for a compilation of *MEFV* pathogenic and likely pathogenic mutations, which we based on the Invefers database (*7*) (including only variants with validated status) and by manually checking the relevant literature. Screening of the present-day Armenian population confirmed the high prevalence of *MEFV* mutations in the region (Table 1; Data S3). In total, we found 13 *MEFV* mutations, with different degrees of penetrance associated with FMF. Consistent with previous studies of regional populations, the four most frequent clinically impactful (and thus routinely tested) *MEFV* mutations in our modern dataset are V726A (7.14%), followed by E148Q (5.77%), M680I(G/C) (3.30%), and M694V (2.75%) (see Table 1). V726A, M680I(G/C), and M694V are predominantly observed in Eastern Mediterranean populations (*14*) but differ in severity: V726A is typically associated with a milder disease course (*15*) but both M680I(G/C) and M694V are associated with more severe phenotypes, with M694V in particular linked to amyloidosis (*16*). In contrast, E148Q is of less certain pathogenicity and is found in healthy individuals across many populations, particularly in East Asia (*17, 18*). Recent evidence, however, suggests it may be pathogenic in specific genetic or ethnic contexts (*19, 20*). A fifth variant, R202Q has the highest frequency in our sample (14.84%), yet it remains of uncertain pathogenicity; a few recent studies suggest it may contribute to FMF symptoms, particularly in the homozygous state, and call for its inclusion in routine genetic screening of patients (*21, 22*). Additional rare impactful and routinely tested FMF mutations detected in our dataset are P369S (1.92%), A744S (0.55%), R761H (0.55%), and F479L (0.27%). These five variants are found primarily among Eastern Mediterranean populations and are associated with variable clinical FMF presentations (*23, 24*). We identified compound heterozygotes and cis complex alleles in the modern Armenian dataset, involving both mild and severe disease-associated mutations. In total, at least one clinically impactful *MEFV* mutation was detected in 41% of the population (58% if R202Q is included).

**Table 1.** Frequency of *MEFV* mutations observed in 182 individuals from the modern Armenian population. Wald confidence intervals were used to calculate the 95% range. *Lower bound was negative and truncated to 0.

| Mutation | Allele frequency (%) | Pathogenicity status by Infevers |
| --- | --- | --- |
| R202Q | 14.84 (9.67-20.00) | Likely benign |
| V726A | 7.14 (3.41-10.87) | Pathogenic |
| E148Q | 5.77 (2.38-9.16) | Uncertain significance |
| M680I(G/C) | 3.30 (0.71-5.89) | Pathogenic |
| M694V | 2.75 (0.37-5.12) | Pathogenic |
| R408Q | 1.92 (0-3.90*) | Uncertain significance |
| P369S | 1.92 (0-3.90*) | Likely benign |
| A744S | 0.55 (0-1.62*) | Likely benign |
| R761H | 0.55 (0-1.62*) | Pathogenic |
| F479L | 0.27 (0-1.03*) | Pathogenic |
| E167D | 0.27 (0-1.03*) | Uncertain significance |
| L110P | 0.27 (0-1.03*) | Likely benign |
| T267I | 0.27 (0-1.03*) | Uncertain significance |

For 85 ancient samples, spanning a time range from ca. 4,530 BCE to 1,600 CE (Fig. 1A), targeted hybridisation capture of all ten exonic regions of the *MEFV* gene (~9.5 kb) was performed, resulting in a mean coverage of 22x across our compiled set of *MEFV* pathogenic and likely pathogenic variant positions. Of the initial 85 individuals, 70 were retained for downstream analyses: 5 samples failed capture, 5 first-degree relatives were removed following kinship analysis, and 5 duplicated pairs were merged. Of the four most frequent, clinically impactful mutations identified in our modern dataset, only E148Q and M694V were recovered among the ancient individuals (Fig. 1B); V726A and M680I(G/C) were not detected in the ancient cohort. We found that a Medieval individual carried the M694V mutation; the high penetrance of this variant (*23*) suggests that this individual living over a millennium ago, may have suffered from the disease. One sample from a site dating to Antiquity carried the E148Q mutation, while another from the same site carried F479L. The most ancient sample from the region with a *MEFV* clinically impactful mutation dates to the Chalcolithic (ca. 4,508 BCE), and carried the P369S mutation. Interestingly, this sample also carried the R408Q variant, which is frequently encountered together with P369S as a cis complex allele (*25, 26*) (Data S4), and a similar case was detected in our modern dataset (Data S3). The P369S variant was also detected in a Late Bronze Age Armenian sample, suggesting a notable prehistoric presence of this variant in the region. In our dataset, we also identified several ancient individuals carrying the R202Q mutation, including one homozygous Bronze Age sample (ca 1,502 BCE).

We examined *MEFV* variant statistics in modern and ancient datasets from the region, restricting comparisons to transversion sites to minimise confounding from postmortem cytosine deamination characteristic of aDNA. Within this filtered set, the ancient dataset showed a higher proportion of variants (including missense variants; fig. S1), thus suggesting a greater genetic diversity in the past. Additionally, there is an excess of novel variants in the ancient cohort (Fisher’s exact test, OR=0.012, 95% CI:0.0003-0.090, p=2.86 × 10^−9^), which likely reflects the underrepresentation of ancient and regionally specific Armenian variation in current reference databases (fig. S2).

To assess the broader geographic and temporal context, we examined 583 published ancient DNA samples (with >1.8x shotgun sequencing coverage) from other regions for the presence of *MEFV* mutations (Data S5). We detected the E148Q variant as early as ~33,000 years ago in western Russia, supporting previous conclusions regarding its ancient origin (*27*) and in later samples from Siberia, Central Europe, Asia, Anatolia and a historical sample for the Andaman Islands. Among other *MEFV* clinically-assayed mutations that we detected were P369S, A744S, M694I, and K695R. We note that some of the above are transition mutations, which are prone to damage in aDNA (*28*) and therefore the results from samples where damage was not eliminated during library preparation must be interpreted cautiously (see Data S5 for a detailed assessment of each sample). We also detected ancient individuals with cis complex alleles carrying E148Q-L110P and P369S-R408Q in Neolithic and Bronze Age Russia, respectively.

### Selection signals for *MEFV* mutations

We investigated evidence of selection on *MEFV* mutations and whether this could be consistent with the well-documented historical dates of bubonic plague pandemics in the region as suggested by Park et al. (*12*). We first conducted a frequency-based between-group association test of clinically impactful *MEFV* variants with allele frequencies greater than 2% in modern Armenians, based on genotype likelihoods (Data S6). We grouped our samples according to their age into pre-first plague pandemic (prior to 541 CE), medieval period, and present-day Armenian cohorts. Three mutations, M694V, M680I (G/C) and V726A, showed significant frequency differences between the pre-plague and modern groups, with V726A also showing a significant difference between the medieval and modern populations. These mutations reach appreciable frequencies in the modern Armenian population (Table 1), but are largely undetected in our ancient dataset from the region (Fig. 2A): M680I(G/C) and V726A were entirely absent, and M694V was found in only a single Medieval sample.

**Figure 2.**
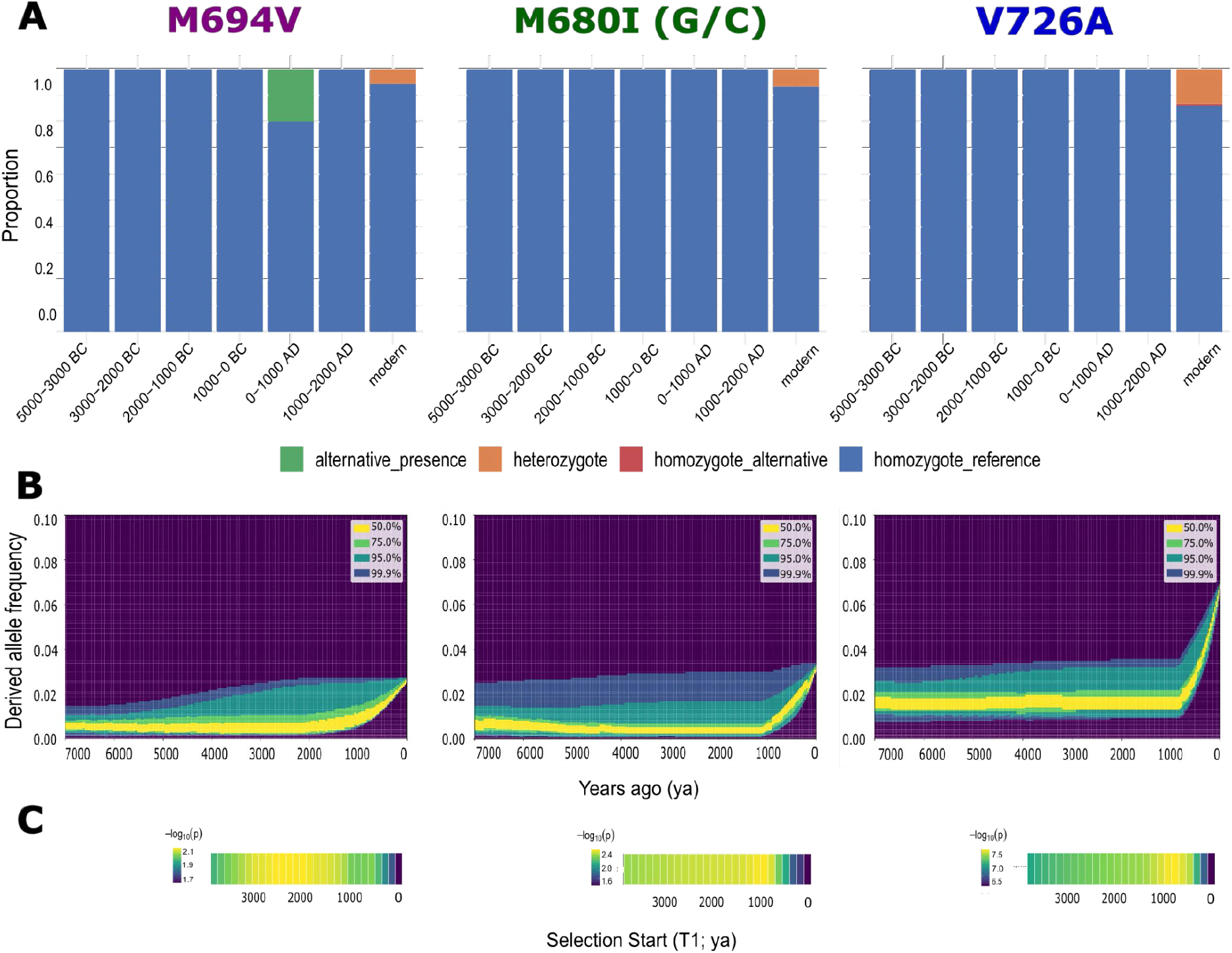
Allele trajectories and selection analyses for the M694V, M680I (G/C), and V726A mutations. **(A)** Frequency bins for carrier status for each mutation over time using combined ancient and modern samples. Only samples with at least two reads supporting each allele were considered. **(B)** Posterior distribution of the inferred allelic trajectory for the best-fitting two-epoch models for *MEFV* mutations calculated using CLUES2 (76 generations for the onset of the second epoch for M694V (S=0.05), 41 generations - for M680I (S=0.09), and 31 - for V726A (S=0.09). **(C)** −log10 P-value surface of CLUES2 two-epoch selection models. M694V, M680I, and V726A were inferred to fit best with two-epoch models, with T1 showing the onset of the second epoch.

We next applied CLUES2 (*29*) to infer the temporal frequency trajectories of variants V726A, E148Q, M680I(G/C), and M694V (the four clinically impactful mutations which showed present-day Armenian allele frequencies greater than 2%) and to formally test for positive selection. We conditioned on the modern Armenian population allele frequencies and evaluated models under constant selection and with either one or two shifts in the selection regime over time. We performed the analyses incorporating ancient genotype likelihoods from our regional time series alongside present-day genealogies inferred with RELATE (*30*). As CLUES2 relies on population size histories to account for the confounding effects of genetic drift when inferring selection, we conditioned our analyses on the Armenian effective population size history inferred by RELATE (fig. S3), which is consistent with previous estimates based on smaller dataset (*10*). We used a grid search approach to infer the onset of selection and its coefficient (*31*), and loci showing significant selection models were identified after correction for multiple testing. In line with the association tests above, CLUES2 identified positive selection for the three pathogenic mutations M694V, M680I (G/C) and V726A (Fig. 2B, Fig. 2C, fig. S4, fig. S5) but not for the less clinically impactful variant E148Q (fig. S5). We found that two-epoch models with sharp recent increases in allele frequencies provided the best fit to the data. For these three variants, positive selection is inferred within recent history with an onset of ~1,000 years or even earlier, with exceptionally large estimated selection coefficients (s=0.09 for both M680I and V726A, s=0.05 for M694V; Fig. 2C; fig. S5). This temporal window is broadly consistent with a scenario in which selection onset may have been driven by plague exposure following the first historically recorded pandemic (541 CE to 767 CE) in the region and later impacted by the second pandemic (1346-1875 CE).

### Haplotype structure and evolutionary history of *MEFV* mutations

We reconstructed the haplotype structure of the abovementioned four clinically impactful *MEFV* mutations (V726A, E148Q, M680I (G/C), and M694V) using RELATE in the Armenian dataset and, where available, in geographically and ethnically diverse comparative modern populations (fig. S6). The inferred tree sequences indicated a monophyletic origin for haplotypes carrying the M694V, M680I(G/C), and V726A mutations (Fig. 3A-C), and further visualisation revealed extended shared haplotype blocks around both mutations (fig. S7-9). Inferred trees also indicated a monophyletic but substantially deeper coalescence for haplotypes carrying E148Q, suggesting a more ancient origin for this mutation (Fig. 3D). However, it should be noted that, as RELATE assumes neutrality, coalescence time estimates for variants under selection can be biased upward, potentially overestimating the true age of mutations (*30*). Notably, E148Q haplotypes from South and East Asian populations formed a tight cluster, consistent with a founder effect that may underlie the high E148Q carrier rates observed in these populations (*32*) (Fig. 3D; fig. S10). Thus, the age difference between the variants mirrors their pathogenicity, with the older and milder E148Q variant reaching high frequency through drift or very weak (undetectable in our study) selection, whereas the younger but more clinically impactful M694V, M680I(G/C) and V726A variants reach comparable frequencies through positive selection.

**Figure 3.**
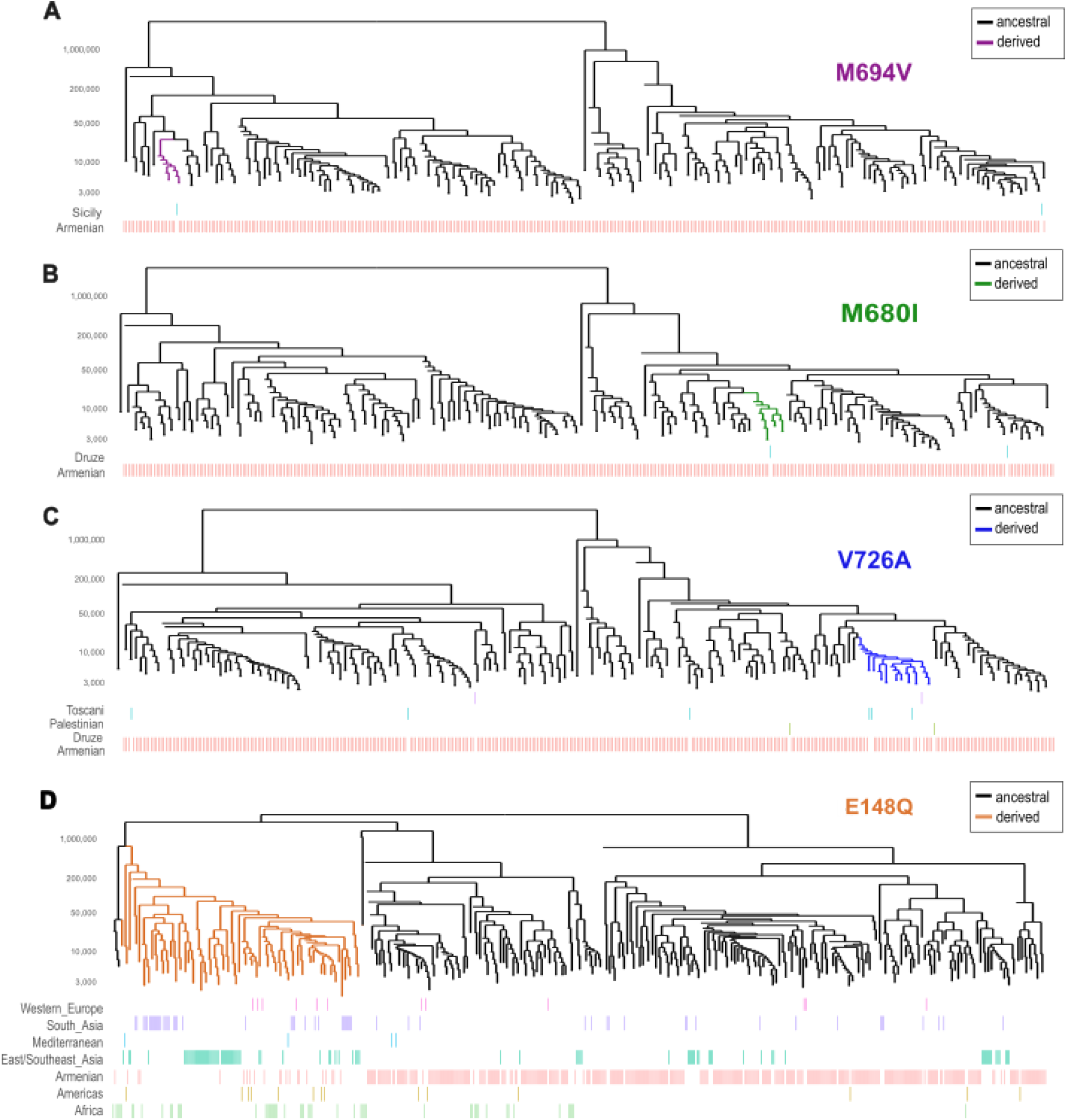
Inferred local genealogies for the *MEFV* mutations M694V (A), M680I (G/C) (B), V726A (C), and E148Q (D) in the modern Armenian and comparative populations. The tree is displayed down to 3,000 generations for clarity; branches are not drawn below this threshold, and their apparent endpoints are a plotting artifact, not biological coalescence events.

## DISCUSSION

Allison’s description of sickle cell heterozygote advantage under malarial challenge over seventy years ago was a major landmark in epidemiology and evolutionary genetics (*33*). Not only was it the first discovery of natural selection in humans but, to great surprise, it showed that the target was a Mendelian disease-causing variant and that the selective force was infection. Apart from this and other malaria-linked examples, *MEFV* is the only other human disease gene which is a likely candidate for having a similar history of strong selection. Accordingly, from extended haplotype analysis in modern Turkish samples, Park et al. (*12*) have argued that its M694V and V726A mutations have been under positive selection and, using functional analyses they identify mitigation of *Y. pestis* infection as a plausible selective agent.

Here we combine ancient and modern genomic evidence to reinforce that the very high incidence of FMF in Eastern Mediterranean populations is due to selection and not the alternate hypothesis of localised drift. First, we find a very high incidence of combined pathogenic mutations of 41% in the modern Armenian population, a world maximum. Second, our time-series approach directly capturing *MEFV* allele frequency trajectories over the past ~6,500 years in this region, which has strong population continuity (*10*), finds that the increases in the three most common Armenian pathogenic variants, M694V, M680I (G/C) and V726A, are both recent and precipitous. Third, our modelling confirms selection as the reason for this increase and the inferred selection coefficients for all the three variants are exceptionally large compared to previously published estimates for other known genetic variants under selection. For comparison, the strongest selection signals in European populations are for the lactase persistence-associated MCM6 variant, estimated in the range of 0.016-0.08 (*34, 35, 36, 37*), and for SLC45A2 (skin pigmentation gene) and C2 (complement immune gene; associated with immune response variation, including infection susceptibility, and psoriasis risk), estimated as 0.02 and 0.04, respectively (*35*).

Human selection coefficients are typically low due to historically small effective population sizes, long generation time, reduced reproductive variance and cultural buffering. Our exceptionally high selection coefficient estimates (comparable in magnitude to those for malaria-linked variants (approx. S=0.12 (*38, 39*)), together with experimental observations that *MEFV* mutations confer an increase in resistance to *Y. pestis* (*12*), infer that bubonic plague exerted one of the strongest selective pressures detectable in the human genome. Our estimates of the timing of selection on all three variants are consistent with an onset linked to the first plague pandemic (or just before) (circa 541 CE-767 CE) and continued selection during subsequent centuries, especially during the second plague pandemic (circa 1346-1875 CE).

Given this linkage and the wide historical impacts of *Y. pesti*s plague, it is striking that *MEFV* variants’ elevated frequency is restricted to Eastern Mediterranean populations. One explanation is that this is the region which alone segregated the variants which were thus available to be selected (*12*). Indeed, each of these resides within a haplotype with a monophyletic origin and has a geography implying an Eastern Mediterranean origin. However, our results, combined with those of Park et al (*12*), extend the evidence for strong concurrent selection to three separate variants each embedded within a distinct haplotype. An argument that the foundation effect of impactful mutation occurred uniquely in the Eastern Mediterranean thrice over seems strained. It is more likely that the geography of these selective effects, as for co-segregating variants conferring malarial resistance (*38, 39*), is driven by an exceptional level of disease challenge.

Bubonic plague manifested in repeated waves, primarily within two centuries-long pandemics: the first pandemic, starting with the Justinian Plague (circa 541 CE-767 CE) and the second pandemic, starting with the Black Death (circa 1346-1875 CE). However, exposure was not even; e.g., historical sources indicate that the Eastern Mediterranean region suffered many more outbreaks during the successive waves of the Justinian Plague; Syria suffered nineteen outbreaks compared to only four in the Balkans, with Italy being similar (*40*). Importantly, the Eastern Mediterranean was likely a permanent enzootic centre with continuous recycling of outbreaks, whereas European epidemics tended to flare and recede (*41*).

That high FMF in the Eastern Mediterranean region is driven by higher selection pressure rather than serendipitous mutations is reinforced by a wider survey of published ancient genomes. We directly observe clinically relevant *MEFV* mutations across a wide geographical range in Eurasia and as early as 33,000 years ago, but none of these reached high frequencies following the emergence of the plague.

## MATERIALS AND METHODS

### Sampling of modern individuals and sequencing

Blood samples were collected from 146 individuals representing the general Armenian population. All individuals were informed about the aim of this study and provided their consent to participate. We interviewed donors about their origins and selected unrelated individuals whose four grandparents were all from the same region within the Armenian highlands (Data S2). DNA samples were extracted at the Institute of Molecular Biology using a modification of the salting-out procedure (*42*). DNA was sent for high coverage (~30x) paired-end sequencing on the Illumina HiSeq X Ten platform (Macrogen, Seoul, South Korea).

### Ethics statements

This study was approved by the Ethics Committee of the Institute of Molecular Biology, National Academy of Sciences of the Republic of Armenia (IRB #00004079).

### Ancient dataset

Ancient samples used in this study come from 21 excavated sites across the region. Detailed description of the sites are provided in Data S1.

### Carbon dating

To confirm archaeological ages of the samples, 7 skeletons from different sites were radiocarbon-dated at Oxford Radiocarbon Accelerator Unit. Mean calibrated ages (BC) with 95.4% low and high probability intervals were obtained by OxCal (Version 4.2; Oxford Radiocarbon Accelerator Unit).

### Ancient DNA extraction and library preparation for initial screening

Petrous bones were preferentially sampled alongside tooth roots. Sample processing took place in a dedicated ancient DNA laboratory at Trinity College Dublin. A wedge of petrous bone was drilled and subsequently powdered using a Mixer Mill. In the case of the tooth sampling, the root part was chopped and powdered. DNA extraction was carried out following various protocols with some modifications (*43, 44*). DNA extraction of ~150 mg bone powder began with incubation in 0.5% bleach for 15 minutes in a termo rotor to remove surface contaminants. The bleach supernatant was discarded, and the remaining pellet was washed three times with UV-ed water. The pellet was then incubated in 1 ml of UV-ed EDTA (0.5M Ph 8) at 37°C for 30 minutes in a constant motion in a thermo rotor. The suspension was centrifuged at 17,000 xg for 1 minute, and the supernatant was collected and stored. The remaining pellet was incubated overnight at 37°C in a thermo rotor, in 1 ml UV-ed extraction buffer (0.47 M EDTA pH 8.0, 0.02M Tris-HCl, 1,7% N-Laurylsarcosine) with added 0.65U of proteinase K following UV. This was centrifuged at 17,000 xg for 10 minutes, and 1 ml of supernatant was transferred to an Amicon Ultra-4 Centrifugal Filter unit 30kDa, with 3mL 10 mM Tris-EDTA buffer and centrifuged at 800xg until 100uL of solution remained. This volume was then added to a silica column (MinElute PCR purification kit, Qiagen, Hilden, Germany) and purified according to the manufacturer’s instructions. DNA was eluted in 50 μL of EB buffer supplemented with Tween 20 (Sigma-Aldrich) (*45*).

16 µL of DNA extract was treated with 3 μL USER enzyme (1 U/µL) and incubated for 1 hour at 37°C to reduce post-mortem deamination lesions. Double-stranded libraries were subsequently prepared for Illumina sequencing following Meyer and Kircher (*46*), with modifications as in Gamba et al (*47*). Libraries followed one PCR amplification (12-14 cycles) using AccuPrime Pfx SuperMix (Life Technologies) with sample-specific 7-nucleotide index primers. Paired-end sequencing (S4, 200 cycles) for library screening was carried out on NovaSeq 6000 platforms by TrinSeq in St James’s Hospital, Dublin, Ireland.

### Ancient sequenced screening data processing

Following sequencing, adapter sequences and low-quality bases at both ends were trimmed from all reads using AdapterRemoval v1.5.2 (*48*), retaining only sequences with a minimum length of 34 bp. Trimmed reads were mapped to the human reference genome (GRCh37/hg19) using BWA v0.6.2 (*49*) with the seed disabled to improve sensitivity for short ancient DNA fragments. Mapped reads were filtered for a minimum mapping quality of 25 and sorted with Samtools (*50*); duplicate reads were removed at the library level using Picard MarkDuplicates v2.22.1 (*51*). Read depth and coverage statistics were assessed using QualiMap (*52*). Endogenous DNA content (EC) and average read length were calculated for each sample to inform downstream capture pooling strategies.

### Ancient DNA capture

The *MEFV* gene is located on chromosome 16p13.3 and spans approximately 14.6 kb. Initially aiming to cover the entire gene with an additional 2,000 bp flanking regions on both sides, due to repetitive regions, we ended up designing a capture bait set of ~9.5 kb with Arbor Biosciences (Hybridization Capture for Targeted NGS), covering all 10 exonic regions of the gene.

Samples with similar average read lengths were combined into groups of up to 10 libraries. Each group aimed to have a total mass of indexed library of 1,000 ng or more following the Arbor Biosciences protocol. Within each group, each sample was represented taking into consideration their endogenous % so that the endogenous DNA contribution of each sample was the same. To achieve the mass required for each sample, several PCR amplifications of each sample library were performed as described for the screening. The number of sequencing reads per pool was assigned to reach approximately 30x coverage across the exonic regions of the target gene.

Each group of Indexed libraries was desiccated prior to hybridisation capture. Hybridisation capture was performed following the Arbor Biosciences protocol. Briefly, a hybridisation master mix was prepared and incubated with the pooled libraries and blocking oligos in a thermocycler overnight at 55°C. The following day, streptavidin-coated magnetic beads were prepared and used to capture the hybridised library fragments, followed by three rounds of stringent washing at 55°C. Post-capture libraries were amplified using KAPA ReadyMix for 14 cycles, and the PCR product was purified using AMPure XP beads prior to sequencing. Paired-end sequencing was carried out on NovaSeq 6000 platforms (S1 100bp).

### Modern sequenced data processing

Adapter sequences were trimmed from the ends of reads using trimadap (*53*). Trimmed reads were aligned to the GRCh37/hg19 build of the human reference genome using BWA and clonal reads were removed with samblaster (*54*). Duplicate reads have been marked with Picard. Base calibration was conducted with the Genome Analysis Toolkit (GATK) version 4.1.9.0 (*55*) and mapping quality filtering was set to 20 with samtools. Variant calling was performed using HaplotypeCaller in GATK to generate intermediate gVCF files, which were further merged with GenomicsDBImport and used in GenotypeGVCFs for joint genotyping of multiple samples. Variant quality score recalibration was performed by VariantRecalibrator for both SNPs and indels (insertions and deletions) separately, and filtered sites were excluded with SelectVariants.

### Capture-sequenced ancient data processing

Post-capture sequencing data were processed following the same steps described above for the screening data. BAM files were merged at the sample level using Picard MergeSamFiles, and indel realignment was performed using GATK. Terminal base quality rescaling was applied by downgrading the quality scores of the first and last two bases of each read to a PHRED score of 2, to reduce the impact of post-mortem damage at read ends. For frequency calculations in ancient dataset, we used GATK HaplotypeCaller, as described for modern samples. We applied an allele balance ≥0.2, and a requirement that the variant site has at least ≥2 reads supporting the alternative allele.

### Authentication of aDNA

DNA damage parameters that are typical for aDNA were estimated using mapDamage 2.0 with default settings (*56*).

### Kinship analyses

Prior to downstream analyses, identical and first-degree relatedness among individuals in both the ancient and modern datasets were assessed using PLINK (*57*), KING (*58*), and READS (*59*) software. Individuals were flagged as related only when consistently identified across all three methods.

### Association test with ANGSD

Association analyses were performed using ANGSD (*60*). Allele frequencies were first calculated separately for each group at the target variant positions using genotype likelihoods (-GL 1, -doMaf 1, -doMajorMinor 1). Association testing was then performed jointly across groups using the score statistic (-doAsso 1), which operates directly on genotype likelihoods and thus avoids hard genotype calling, making it suitable for low-coverage ancient DNA data.

### Phasing

Phasing of modern samples was conducted with SHAPEIT5common (*61*), without using a reference panel. Because M694V had a low number of haplotypes in the Armenian dataset, we included an additional sample from Sicily during phasing only. Phasing was repeated three times for each variant to confirm consistency of selection analyses per each phasing run.

### Selection

To infer the evolutionary history of *MEFV* variants, phased VCF files were first converted to RELATE-compatible haplotype format using RelateFileFormats (--mode ConvertFromVcf). Input files were then prepared using the PrepareInputFiles script in RELATE v1.1.9 (*30*), which applied a strict accessibility genome mask to exclude low-confidence regions and used the Ensembl human ancestral genome sequence (human_ancestor_GRCh37, e59) to orient alleles to their ancestral state. Genealogical trees were subsequently reconstructed across all 22 chromosomes using RELATE, applying a mutation rate of 1.25×10^−8^ per base pair per generation and an effective population size of 30,000. Population size changes through time were then estimated per chromosome and subsequently re-estimated jointly across all 22 chromosomes using the EstimatePopulationSize script, with 10 iterations and a generation time of 28 years, to provide a cohesive coalescent framework for downstream selection inference.

For selection inference, local genealogical trees were extracted for each target *MEFV* variant using RelateExtract, and branch lengths were resampled 500 times per variant using the SampleBranchLengths script. Derived allele frequencies at each target position were obtained from ANGSD-based allele frequency estimates in the modern Armenian dataset, ensuring correct ancestral/derived allele orientation. Resampled trees were converted to CLUES2 (*29*) input format using the RelateToCLUES.py script.

In order to find the time of the change in selection regime (T1), we performed a grid search as described in (*31*). We first tested two-epoch models, varying T1 between 6 and 141 generations ago in steps of 5 generations. We then tested three-epoch models, varying T1 between 51 and 141 generations and T2 between 6 and 46 generations, using the same step size. P-values were corrected for multiple testing using Holm’s sequential Bonferroni procedure. The analyses have been repeated three times per each phasing run to confirm consistency of results.

### Haplotype visualisation

We merged a comparative dataset from 1000 genomes and HGDP panel (*62*), proceeding with phasing and RELATE steps as described above. Genealogical tree visualisations for individual variants were generated using RELATE’s TreeViewMutation.sh script. Haplotype structure was visualised using HaploStrips (*63*).

### Annotation

Variants for both the ancient and modern cohorts were annotated using the Ensembl Variant Effect Predictor (VEP). Sites within the ancient cohort were filtered to retain transversions only, applying an allele balance ≥0.2, and a requirement that at least one sample per site show ≥2 reads supporting the alternative allele. The resulting set of ancient variant positions was then used to extract corresponding sites in the modern cohort, enabling a direct, site-matched comparison of variant consequence spectra between the two cohorts.

## Funding

EU MSCA-IF under grant agreement 101063265 (A.H.)

European Research Council grant agreement 885729-AncestralWeave (D.G.B. and A.H.) Science Committee of the Ministry of Education and Science of Armenia (research project no. 21AG-1F025) (L.Y. and Z.K.)

## Author contributions

A.M., D.G.B., L.Y., and A. H. conceived and designed the study. A.M. and D.G.B. supervised the study. A.H. conducted bioinformatics analyses. L.M.C., I.J., and E.B. contributed to the data analyses. Laboratory work was done by A.H., V.M., B.A., and Z.K. All other authors gave access to samples, provided archaeological and osteological context and aided in the interpretation of results. Funding for modern whole-genome generation was acquired by L.Y. and H.S. The manuscript was written by A.H., A.M., and D.G.B., with contributions from all co-authors.

## Competing interests

All other authors declare they have no competing interests.

## Data, code, and materials availability

All data and code needed to evaluate and reproduce the results in the paper are present in the paper and/or the Supplementary Materials. The generated raw Fastq files covering the *MEFV* gene will be available upon publication through the European Nucleotide Archive (ENA).

## Supplementary Materials

Supplementary Text

Figs. S1 to S10

Legends for Supplementary Tables

## Notes

### Competing Interest Statement

The authors have declared no competing interest.

### Summary of Updates

Uploaded current pdf file of the main manuscript

