## Supplementary Material for "Plague-driven selection enriched familial Mediterranean fever mutations in Armenia"

Supplementary Text

**Description of the archaeological sites**

**Agarak-2:** The Agarak 2 burials were excavated in the northwestern section of the Agarak Historical-Cultural Reserve, situated on the western edge of Agarak village in the Aragatsotn Province of the Republic of Armenia, on the bank of the Amberd River. The burials were discovered within a multi-room medieval residential house. A total of 29 burials were uncovered inside the rooms of this early medieval structure, performed after the building had lost its original function and was abandoned.

The excavated graves primarily belong to two structural types: cist graves and pit graves. Both cist and pit graves are oriented along an east-west axis, with the side walls of cist chambers built of unworked stones of various sizes and covered with stone slabs. Pit burials were interred in wall openings, directly on the bedrock after removing the clay-plastered floor, on or within the clay floor, or in higher layers above it. The deceased were generally placed in a supine position (on their backs), with hands placed on the abdomen, crossed over the chest, or on the pelvis, adhering to Christian funerary customs. The grave goods were virtually absent, with only a needle and an ornamental pin recovered from cist burials on the left side of the rock platform.

Stratigraphic observations, architectural contexts, and radiocarbon dates indicate three main chronological phases for these Christian burials:

• 6th century AD: Pit burials interred within wall openings that were later sealed during structural repairs (Burials No. 3, 4).

• 7th–8th centuries AD: Cist and pit burials performed directly on the bedrock immediately following the abandonment of the building (cist: Burials No. 5, 9, 12, 15, 16, 17, 19, 22; pit: Burials No. 1, 18, 27).

• 12th–13th centuries AD: Cist and pit burials executed on or into the clay floor or higher fill layers (cist: Burials No. 6, 8, 14, 23, 24; pit: Burials No. 10, 11, 20, 21, 28, 30).

**Aghtsk:** The royal tomb of Aghtsk is located in the Aghtsk community of the Aragatsotn Province of the Republic of Armenia at an elevation of 1,230 m asl. The monument has been known since as early as the 4th–5th centuries AD in written sources. This multi-layered site was excavated periodically between 2015 and 2022. In the central part of the site, a medieval cemetery was uncovered. So far, the excavated remains are around 256 individuals from the High and Late Middle Ages, of which 185 have been identified (18 newborns, 61 children, 47 females, 43 males, and 16 undetermined).

The burials followed Christian funerary practices - bodies were laid on their backs, generally facing east, with extended torsos, hands crossed over the chest or abdomen, and legs placed parallel. The skeletons were found at depths of 30-100 cm below the surface, densely packed in 3-4 layers, often directly on top of one another. They date to the High and Late Medieval periods (*64*).

**Artanish 23:** This Iron Age necropolis in the northwestern Artanish Peninsula (1,927 m asl, Gegharkunik Province) was first identified by Yervand Lalayan in the early 20th century and surveyed by an Armenian-German expedition in 2015-2016. It holds more than 30 visible cromlechs and four large cromlech-ringed mounds; ten tombs were excavated in 2019-2022. Tomb No. 1, round-cromlech and slab-built, runs east-west and held a 30-35-year-old male with vessels, bronze and iron arrowheads, bronze bracelets, rings, and beads documented in situ. Fauna was limited to sheep/goat and cow/bull (Bobokhyan and Kunze, 2021; osteology by N. Zarikyan).

**Artanish 29:** About 1 km from Artanish 23, in the western Artanish Peninsula (1,935 m asl, Gegharkunik Province), this Iron Age necropolis - found by the same 2015 expedition - has more than 20 round-cromlech tombs. In 2019 and 2021, two tombs were excavated. Tomb No. 1 is a burial mound encircled with a round cromlech and has stone-soil armour. А cist grave was opened in the centre of the cromlech with a west-east orientation. The walls of the chamber were lined up quite neatly and canonically. Meanwhile, the biological and archaeological materials inside had no canonical arrangement. The chamber was full of human bones, animal bones (sheep/goat, cow/bull, pig, wolf; the osteological material is identified by N. Zarikyan), ceramic sherds, beads, metal, and bone objects that were buried here simultaneously (*65*).

**Artik st Astvatsatsin church:** The historical core of the modern town of Artik (Shirak Province, Armenia) developed around two early medieval churches: St Astvatsatsin (St Marine), dated to the 5th–7th centuries CE, and St Gevorg (St Lusavorich), dated to the 6th–7th centuries CE. Remains of 19th-century Armenian vernacular dwellings (glkhatun-type houses) and associated structures are also preserved within the area between the two churches. Archaeological excavations directed by Levon Mkrtchyan were conducted in 2019 and 2021 as part of the restoration programme for St Astvatsatsin Church. The investigations documented several phases of construction, renovation, and use, principally dating to the 10th-13th and 19th-20th centuries CE. The 10th-13th-century deposits were particularly substantial and reflected extensive medieval rebuilding and modification of the church. These activities had heavily disturbed or removed much of the archaeological stratigraphy associated with the original 5th-7th-century foundation phase.

Two complete khachkars (cross-stones), stylistically attributable to the 11th-13th centuries CE, were preserved in the southwestern sector of the church. Together with the burials uncovered immediately outside the southern wall, they indicate the presence of a medieval cemetery associated with the church. Several graves extended to the level of the stylobate, an architectural element belonging to the early medieval building phase. Their stratigraphic relationship indicates that, by the time the cemetery was in use, the lower exterior elements of the church had already been covered by accumulated deposits. One of the individuals included in the present genetic study derives from this burial group.

**Bjni:** Located on the right bank of the Hrazdan River, east of the medieval fortress (1,620 m asl). The material for this study comes from Tombs No. 3 and 4, both encircled by cromlechs of large stones and situated northeast of Tomb No. 2. Tomb No. 3, built on a hill with a southeastern slope, has a chamber oriented northwest-southeast. Finds were scattered in the fill, mainly pottery and bones, with concentrations along the northern wall. A black burnished vessel with grooved ornamentation (mid-2nd millennium BCE) contained bones and a bronze dagger fragment. A secondary cromlech with a central standing stone was documented in the northeast. Tomb No. 4, located 10 m northwest of Tomb 3, has a north-south oriented chamber. The northern section contained a black jug, an antler-shaped object, and pottery along the western wall. Animal bones, including two skulls, were placed near the northern wall, while the deceased lay in the southern part of the chamber with the skull oriented westward. A reddish-brown vessel (mid-2nd millennium BCE) was found outside the eastern cromlech. Both tombs share architectural and mortuary traits, including cromlech enclosures, central chambers, and deposits of ceramics, metal objects, animal remains, and human burials, dating to the middle of the 2nd millennium BCE.

**Dvin Eranos:** The Dvin-Yeranos burial ground sits on a plateau 1,600 m above sea level, on the slope of the Yeranos Mountains massif, 13 km north of Dvin village and bordered by a gorge to the east. First excavated in 2020, the site yielded three tombs from its eastern edge along the road to the gorge, containing ceramics, decayed skeletal remains, jewelry, and other artifacts dating to the Late Bronze-Early Iron Age. Unstudied rectangular structures were also identified within the burial ground, and further research is expected to expand the site's known extent. Some recovered finds are now exhibited at the Erebuni Museum. Evidence indicates the slopes of Mount Yeranos were continuously inhabited from the 10th century BC, with the sheer number of tombs pointing to a long-lived, active local population. Despite terrain unsuited to farming, people continued to live in the area until the first half of the 20th century.

**Hatsarat:** The tomb was uncovered in 2016 during construction work at the Berd Glukh archaeological site, located within the administrative boundaries of the Hatsarat community (1915 m asl). The burial chamber is soil-cut and oriented east to west. At a depth of about 1.15 meters, artefacts dating to the late period of the Kingdom of Van (Urartu), specifically the 7th century BCE, were discovered. These included red polished ceramic vessels, two glass beads, a fragment of a bronze object, and a piece of a bone inlay decorated with a carved motif-likely part of an inlaid item belonging to another object.

**Keren:** Tomb No. 97 is located along the Keren village road (570 m asl). The tomb is a stone cist oriented north-south, deviating about 20 degrees toward east-west. The walls of the stone cist were slightly inclined inward, indicating the presence of a false vault. In the southern part, the wall was constructed not from bedrock but from the level of the earthen “pillow.” The simultaneous uncovering of the stone-earth fill of the chamber and the archaeological material suggests that the finds were placed while the tomb was being filled. The greatest concentration of material was observed in the southern part of the tomb, arranged in several layers and reaching almost to the surface. In the northern part, the finds were more sparsely distributed, but from here metal objects were uncovered: votive plaques, pendant-medallions, rings, tubular beads, as well as fragments of a vessel.

**Kanagegh:** The Kanagegh archaeological site is located on the western shore of Lake Sevan, about 5.5 km north of the village of Yeranos, in the Gegharkunik region of Armenia (1,940 m asl). The site consists of a settlement and a burial field, occupying approximately 200 hectares. In the burial field, 30 burial mounds have been preserved. From 1980 to 2024, eleven of these were excavated. Archaeological finds indicate that the earliest burials in the cemetery date back to the 15th-14th centuries BCE. In some burial chambers, reburials were performed during the Early Iron Age (12th-9th centuries BCE) and the Classical period (4th-1st centuries BCE). Excavations in the Kanagegh burial field have uncovered bronze daggers, diadems, bracelets, a Mitanni cylinder seal, stone moulds for casting ornaments, and numerous luxury items (beads, earrings, necklaces, rings, brooches, etc.).

**Karin Tak:** is located in the midst of the lush Karintak Forest. Karintak cave contains two separate cave passages (termed Cave 1 and Cave 2). Cave 1 is considerably longer, extending for approximately 42 m in north-east direction (inward from the entrance). The passage continues deeper, beyond this point, but is too narrow to pass through. Cave 1 actually narrows and widens (constriction and necking) several times along its length, producing a set of small ‘sub-chambers’. Cave 2 extends in from the entrance in a westerly direction for only c. 10 m before turning 90° in a southerly direction for another c. 5 m.

**Makravank:** is located on the border of Nig and Varazhnunik (Tsaghkunk) provinces of the Ayrarat state, at the foot of the Tsaghkunyats mountain range (1783 m asl). The origin of the name “Makravank” is uncertain; it may have referred to a monastery for chaste hermits or for healing from diseases such as leprosy (Mesrop Smbatyants). Information about the monastery is preserved only in late medieval written sources. The complex, entirely built of polished basalt, consists of a cathedral church with a central dome (early 13ᵗʰ century), an eastern two-story depositary, a ruined courtyard (early 13ᵗʰ century), and a southern single-nave church with a double-sloped roof (10-11ᵗʰ century). The churches were restored in the 1980s, and a medieval cemetery with khachkars and tombstones surrounds the complex. Excavations were conducted in 2018 by the Institute of Archaeology and Ethnography, NAS RA, prior to partial restoration works. The courtyard and the surroundings of the two churches were investigated, yielding numerous artefacts, including pottery, metal objects, khachkars, and tombstones. Inscriptions were also supplemented, providing additional historical context. The findings shed light on the history and development of Makravank (*66*).

**Metsamor:** The Metsamor archaeological site is situated in the Ararat Valley, 35 km southwest of Yerevan, close to the marshy sources of the Metsamor River (850 m asl). The first information referring to this site was recorded in 1890 (*67*), and the first excavations were carried out in 1959-1962 (*68*). The settlement was occupied from the Early Bronze Age (Kura Araxes culture) to medieval times (*69*). The faunal assemblage indicates that cattle, sheep, and goat husbandry were practised at the site. Additionally, horse, donkey, and camel remains were present in the faunal material, together with representatives of the local wild game (red deer, gazelle, beaver, and hare) (*70*).

**Nerkin Naver:** is located 35 km west of Yerevan (1080 m asl) and consists of a kurgan field, caves, and circular stone structures. Excavations were conducted in 2002-2025 by the expedition of the Scientific Center for Research of Historical and Cultural Heritage (dir. H. Simonyan). Among the investigated features, Kurgan No. 4 was notable for its double-layered burial structure. The upper layer, beneath a stone-and-earth mound, contained the burial of a 35-45-year-old male bodyguard, while below, on a prepared tuff slab, lay the severed head of his master. Grave goods included silver earrings, bronze padlocks, glass and faience beads, a decorated clay mug of Beden type, and a glass eye-bead. The lower layer and undercut northern grave yielded multicoloured beads (including carnelian, amber, and gold-plated gypsum), red- and black-polished vessels, a bronze dagger, pin, mirror-standard, basalt piala, seashells, obsidian flakes, and matting remains. Particularly notable were gold-plated gypsum wheels, probably belonging to a model ritual cart. Kurgan No. 4 is dated to the early Middle Bronze Age (ca. mid-3rd millennium BCE) (*71*).

**Odzaberd:** is located on a hill east of the town of Tsovinar, at the southeast corner of Lake Sevan, Gegharkunik Province, Armenia, at 1,921 m asl. Excavations conducted in 2018 (locus H) uncovered a 16-meter-long section of massive walls built with semi-dressed stones, approximately 2 meters wide. Clay floors, an economic well, and structural fences suggest a domestic compound enclosed by an outer fence. Based on stratigraphic observations, the structure likely dates to the post-Urartian period, specifically the late 7th to 5th centuries BCE. Finds included numerous pottery sherds, fragments of portable hearths, metal objects, animal bones, and other domestic refuse. At a later stage, during the Late Classical period, burials were placed along the southwestern outer corner of the wall, extending northward. A collective burial contained five individuals: one child (5-6 years old), two females (14-17 and 35-45 years old), and two males (30-35 and 50-55 years old), mostly in flexed positions, with one skeleton disarticulated. To the north, a single jar burial contained a male (35-45 years old) in a flexed, embryonic position. No grave goods were associated with the jar burial. These burials indicate that by the Late Antique period, the outer city had been partially abandoned and repurposed as a burial ground.

**Shengavit:** Shengavit is an Early Bronze Age site in Armenia, built up in multiple occupation layers to a depth of four meters. The settlement sits on an irregularly oval platform covering six hectares and was enclosed by a substantial stone wall. Beneath a tower attached to this wall, on its northern edge, archaeologists uncovered a stone-slab-covered underground passage that led down to the Hrazdan river, while the site's burial ground stretched out beyond the fortification. Shengavit's architecture reflects a mix of long-standing building customs and clear advances in construction technique. Hakob Simonyan has led excavations at the site since 2000; a joint Armenian-American archaeological expedition, co-headed by Simonyan and Mitchell Rothman, worked there from 2009 to 2012.

**Sonasar:** The mausoleum was discovered in the elevated valley of the Sodk region, approximately 10 km from the village of Sonasar (*72*), in the historical province of Siunik (1,350 m asl). The burial dates to the late 1st century BCE. The tomb structure, along with the associated burial and grave goods, was found largely intact. Among the finds were fragments of burial furniture, including carved and decorated legs, as well as personal ornaments and luxury items made of gold, silver, and bronze. The burial likely belonged to a woman of middle social standing, interred with rich and aesthetically refined objects. Notable finds include a gold necklace with an oval medallion containing green glass, set into a shield-shaped plaque; another necklace with bird-head terminals and a convex agate gem featuring the relief of a vulture; and two gold rings - one bearing an image of Athena standing with an owl, the other with the figure of a standing youth. A pair of earrings depicting a worship scene of the Mother of Gods, Cybele, and her consort Attis, was also recovered. These items show strong parallels with objects from the tomb of Volta Finta in the Sisian region of Siunik. In particular, a silver cup from that tomb bears an Aramaic inscription naming Araqszat, meaning "protected by the gods", a title typically reserved for rulers or high officials. This suggests a shared cultural context between the Sonasar and Sisian tombs. Both mausoleums date to the late 1st century BCE, reflecting the wealth, artistic expression, and funerary customs of Siunik’s elite class during this period.

**Uyts:** Uyts is one of the largest settlements documented in southern Armenia, sprawling across as much as 200 hectares on the south bank of the Vorotan River, just east of Sisian (*73*). The site occupies a position on steep, rocky heights overlooking the surrounding landscape, and its defenses reflect that strategic setting: several well-preserved lines of massive fortification walls, constructed from roughly-worked basalt blocks, still stand as an imposing feature of the site. Traces of structures from multiple occupation phases are visible both within the fortress and in the surrounding area, alongside numerous burials, including kurgans and stone cists, some quite large, that likely span a long chronological range, though most have been disturbed by looting, possibly since antiquity.

Intensive surface survey has shown that Uyts was occupied across an unusually long span of time, from the Early and Middle Bronze Age through the Early Iron Age and into the Classical and Medieval periods. Excavation began in 2006 with two small test trenches. Trench DT-2 uncovered material from the Early Yervandid period, with radiocarbon dates placing this occupation phase in step with the corresponding phase at Shaghat I. Trench AT-1, expanded further in 2007, revealed dwelling remains, burials, and artifacts dating to the Early Iron Age.

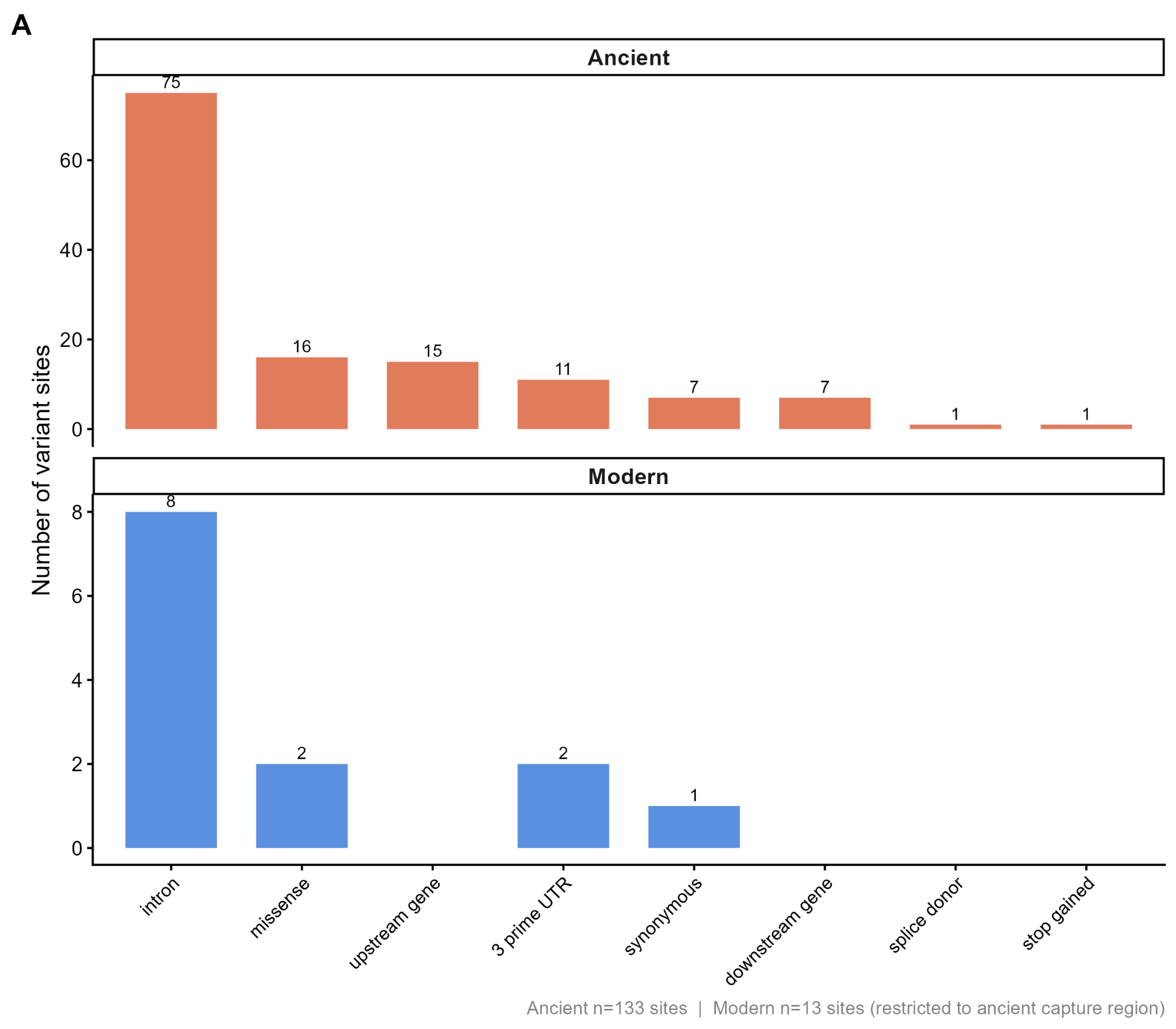

Fig. S1.

**VEP-annotated consequence categories for MEFV variant sites in ancient (n=70) and modern Armenian (n=182) cohorts.** Sites within the ancient cohort were filtered to retain transversions only, applying an allele balance ≥0.2, and a requirement that at least one sample per site show ≥2 reads supporting the alternative allele. The resulting set of ancient variant positions was then used to extract corresponding sites in the modern cohort, enabling a direct, site-matched comparison of variant consequence spectra between the two cohorts.

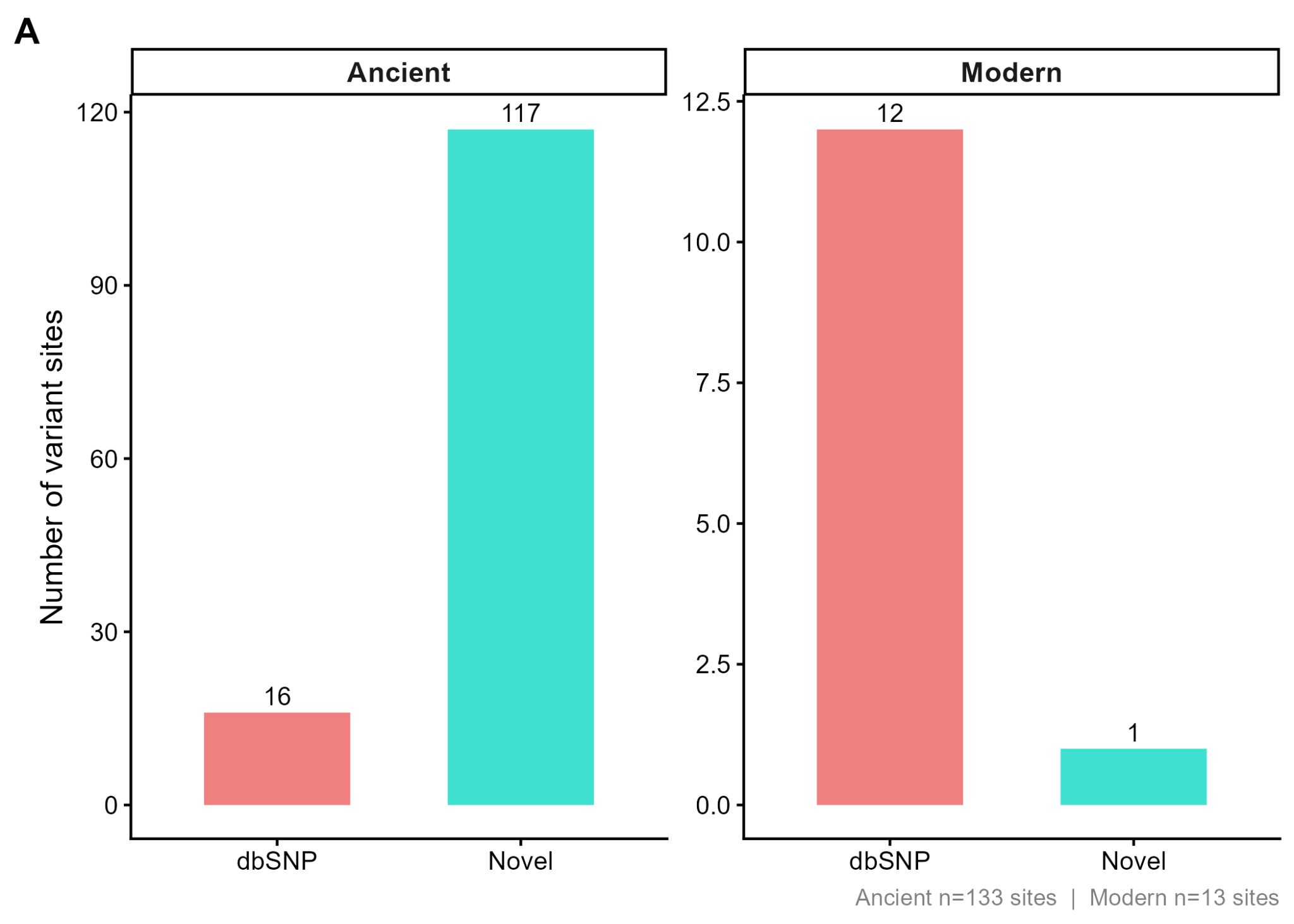

Fig. S2. Number of variant sites classified as known (dbSNP) or novel in ancient (n=70) and modern Armenian (n=182) cohorts. Sites within the ancient cohort were filtered to retain transversions only, applying a minimum per-sample read depth of 2, an allele balance ≥0.2, and a requirement that at least one sample per site show ≥2 reads supporting the alternative allele. The resulting set of ancient variant positions was then used to extract corresponding sites in the modern cohort, enabling a direct, site-matched comparison of variant consequence spectra between the two cohorts.

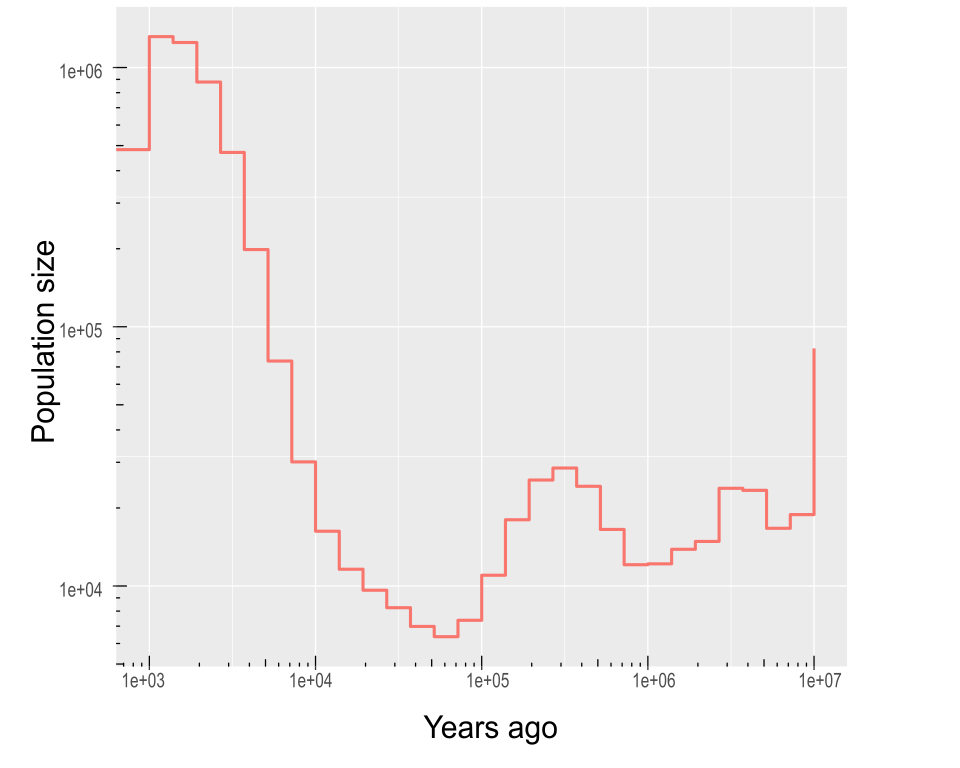

Fig. S3. Estimated effective population size history of the modern Armenian population.

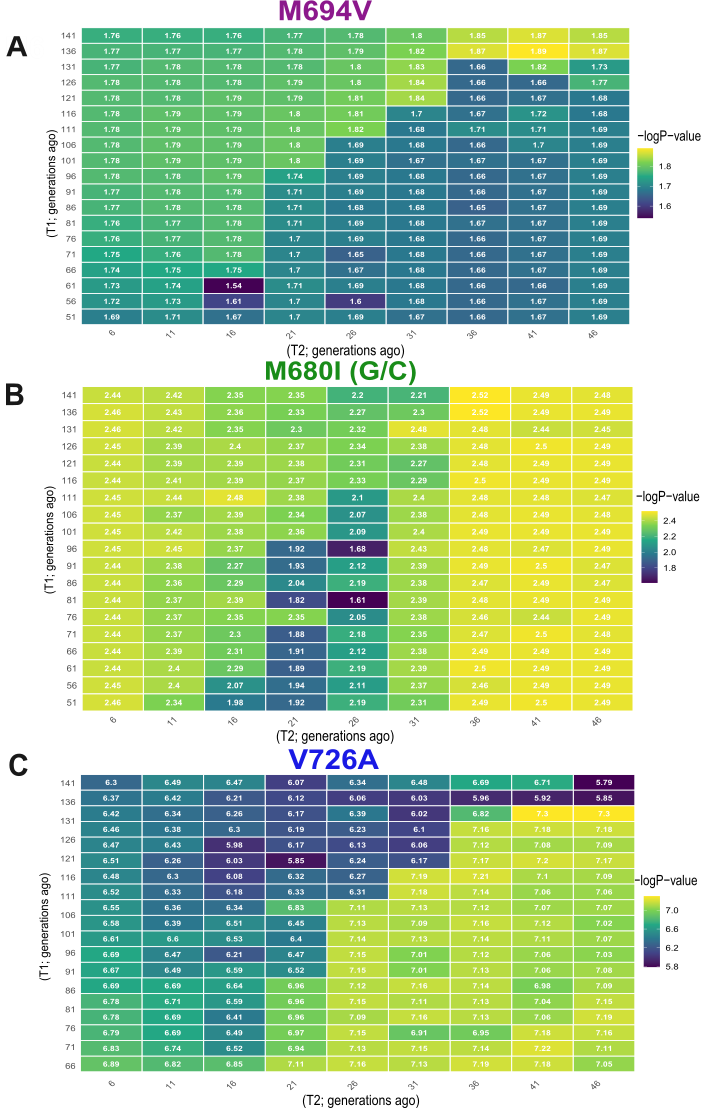

Fig. S4. −log10 P-value surface of CLUES2 three-epoch selection models for M694V (A), M680I (B), and V726A (C) variants.

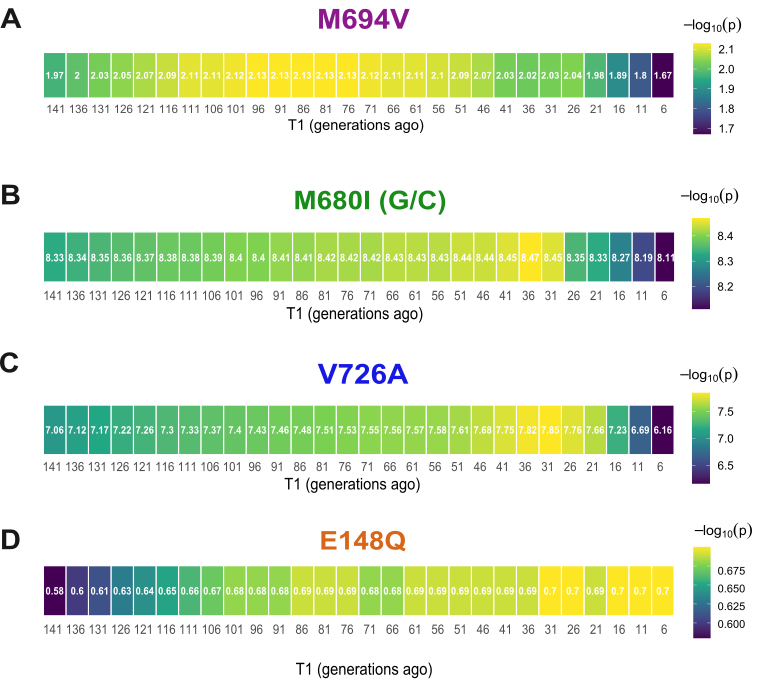

Fig. S5. −log10 P-value surface of CLUES2 two-epoch selection models. M694V (A), M680I (B), V726A (C), and E148Q (D) were inferred to fit best with two-epoch models, with T1 showing the onset of the second epoch.

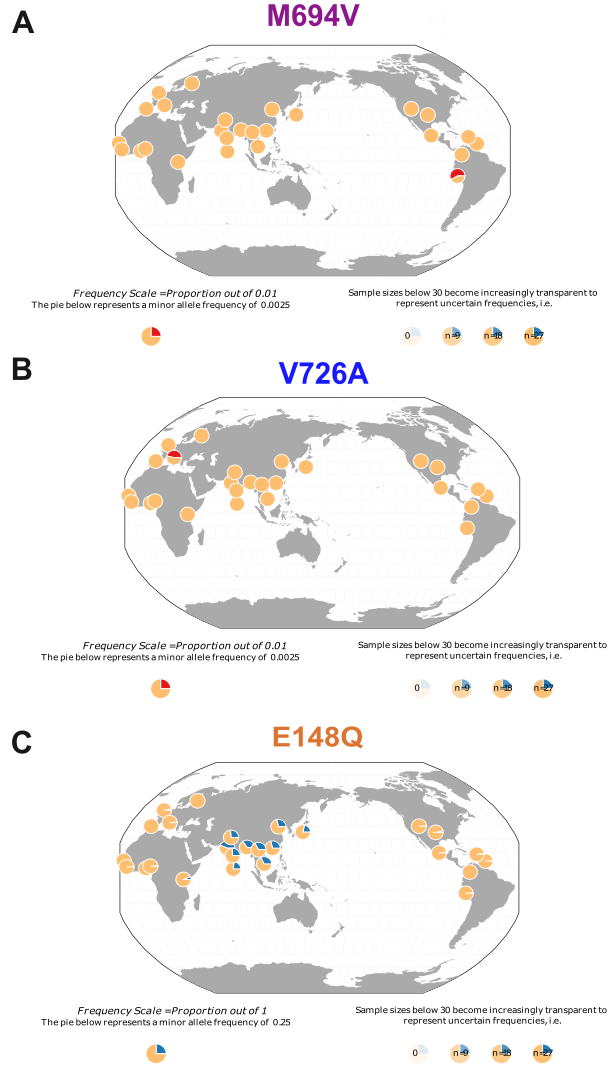

Fig. S6. Frequencies of MEFV mutations M694V (A), V726A (B), and E148Q (C) in the 1000 Genomes database. M680I(G/C) is absent from all populations. Note that a M694V mutation was found in one individual from Peru; this individual was excluded from analyses due to a high level of admixture.

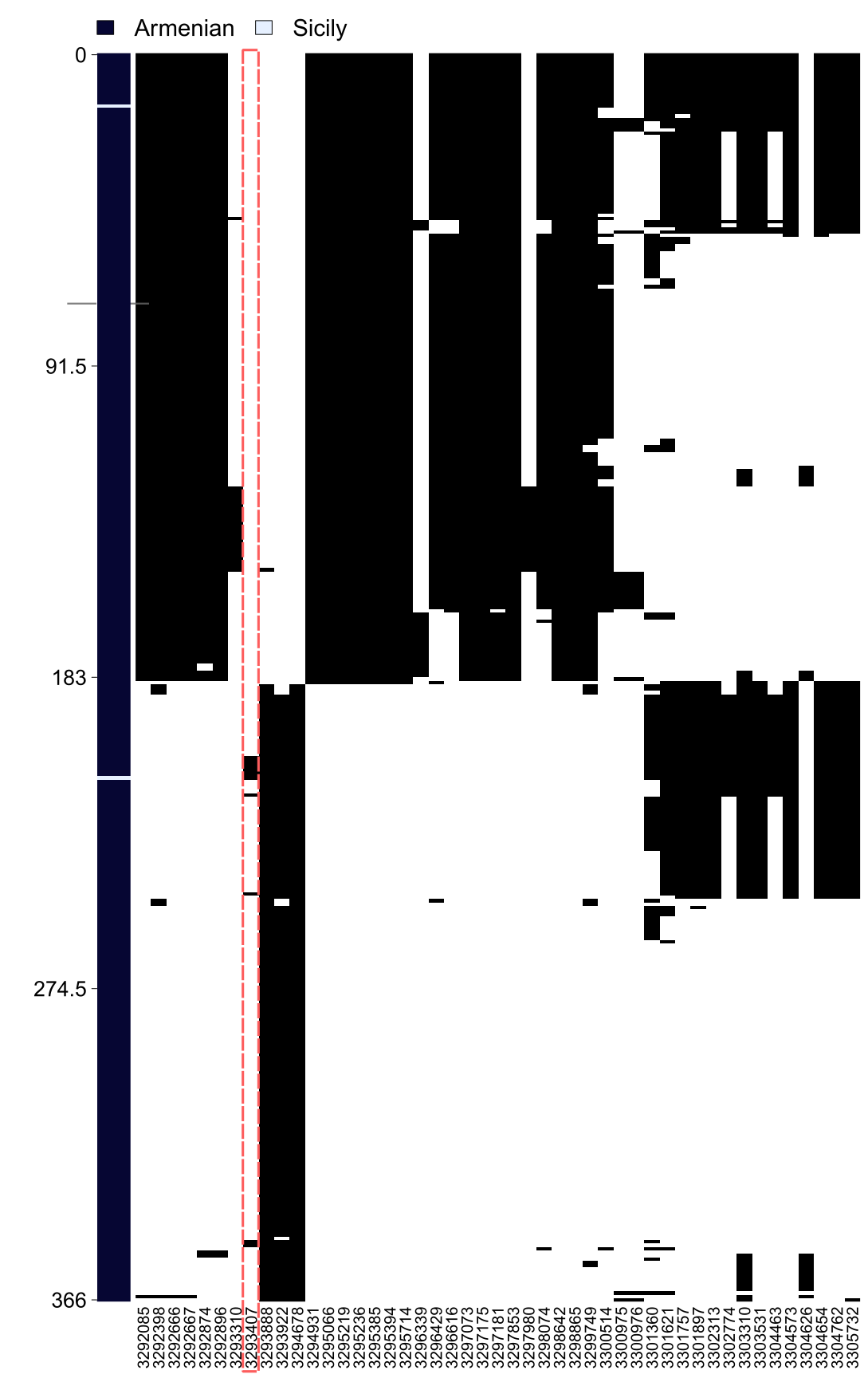

Fig. S7. Haplostrips plot showing haplotype structure surrounding M694V variant in the Armenian and comparative dataset. Each row represents a single haplotype and columns represent SNPs, ordered by physical position; haplotypes are coloured by ancestral/derived allele state.

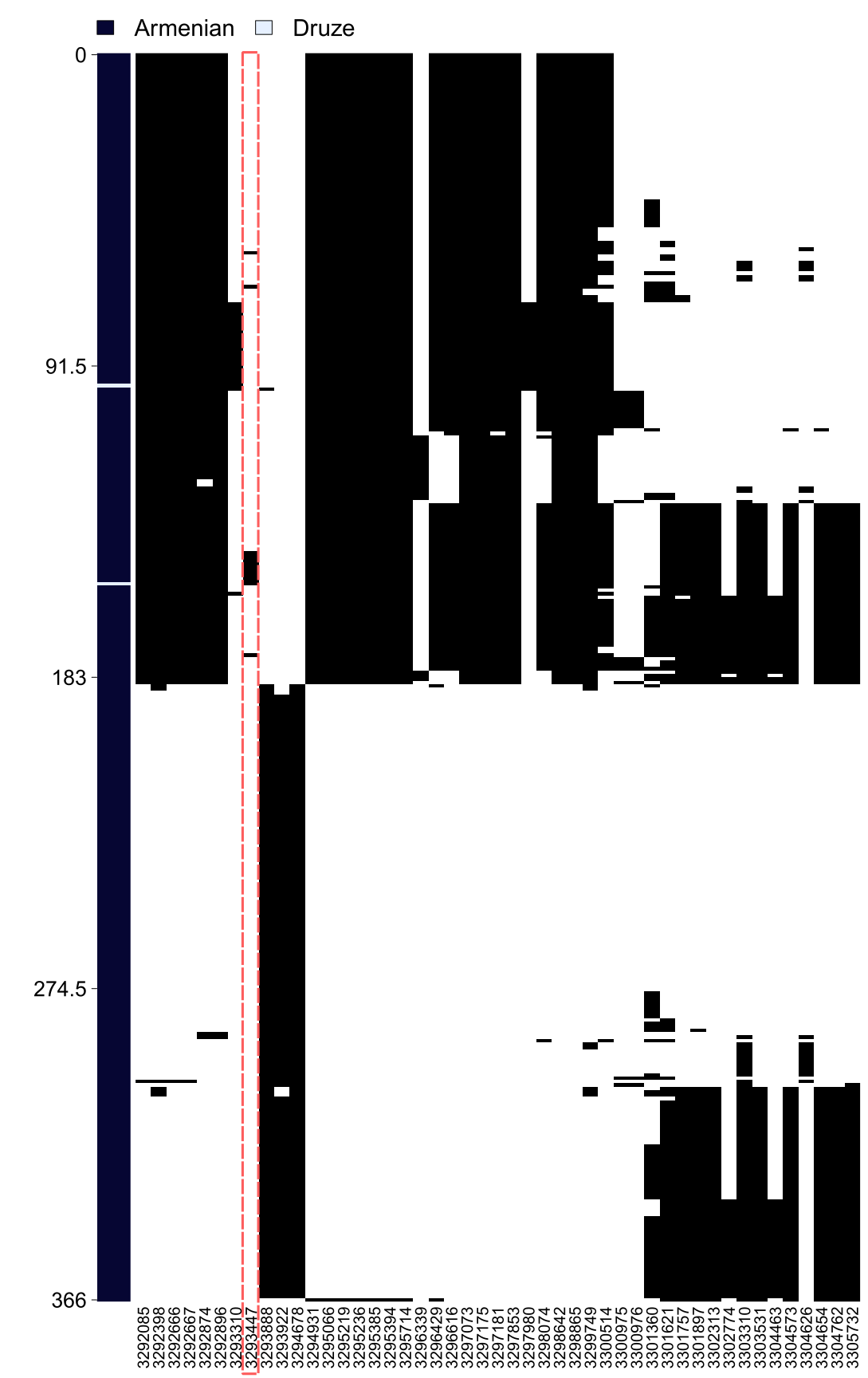

Fig. S8. Haplostrips plot showing haplotype structure surrounding M680I variant in the Armenian and comparative dataset. Each row represents a single haplotype and columns represent SNPs , ordered by physical position; haplotypes are coloured by ancestral/derived allele state.

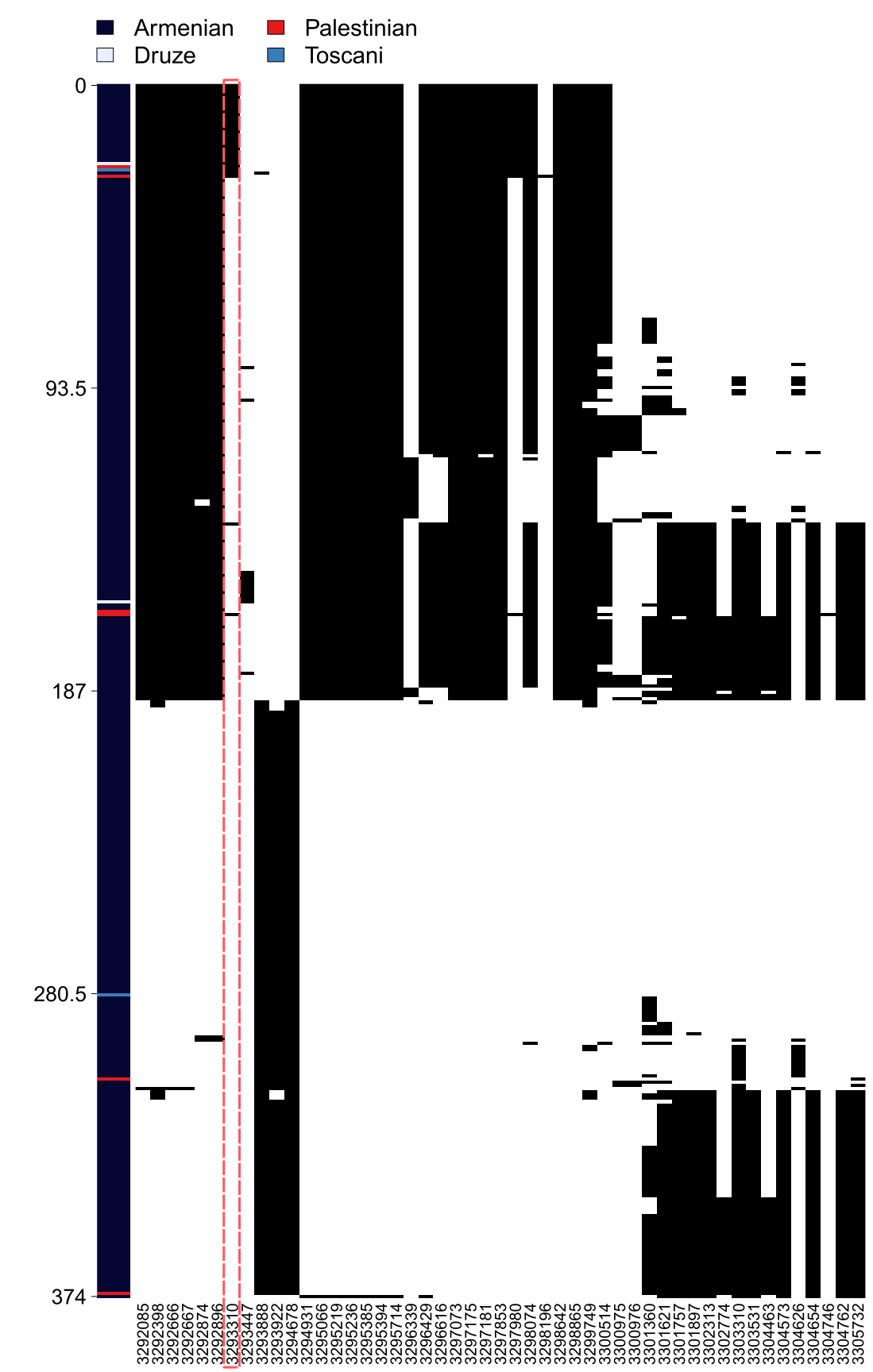

Fig. S9. Haplostrips plot showing haplotype structure surrounding V726A variant in the Armenian and comparative dataset. Each row represents a single haplotype and columns represent SNPs, ordered by physical position; haplotypes are coloured by ancestral/derived allele state.

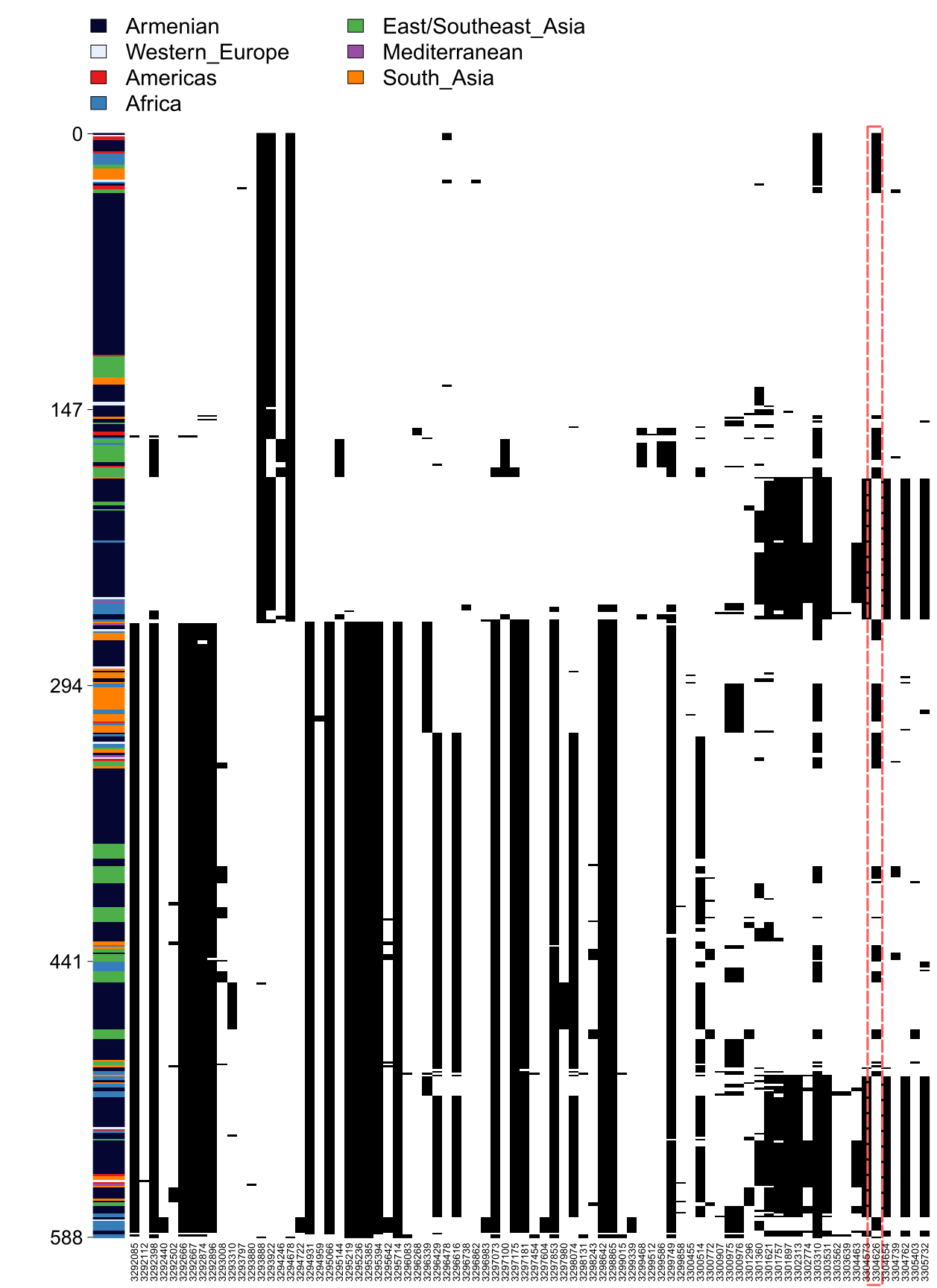

Fig. S10. Haplostrips plot showing haplotype structure surrounding E148Q variant in the Armenian and comparative dataset. Each row represents a single haplotype and columns represent SNPs , ordered by physical position; haplotypes are coloured by ancestral/derived allele state.

Legends for Supplementary Tables

These tables are too large to include within this .pdf file, they are provided as separate files

(.xlsx)

Data S1-6.xlsx

Data S1. Metadata for ancient samples.

Data S2. Metadata for modern Armenian samples

Data S3. Screening results for modern Armenian samples

Data S4. Screening results for ancient samples from Armenia

Data S5. Screening results for comparative ancient samples

Data S6. ANGSD results for mutations passing multiple correction
